# State-dependent effectiveness of cathodal transcranial direct current stimulation on cortical excitability

**DOI:** 10.1101/2023.03.03.531020

**Authors:** Alessandra Vergallito, Erica Varoli, Alberto Pisoni, Giulia Mattavelli, Lilia Del Mauro, Sarah Feroldi, Giuseppe Vallar, Leonor J. Romero Lauro

## Abstract

The extensive use of transcranial direct current stimulation (tDCS) in experimental and clinical settings does not correspond to an in-depth understanding of its underlying neurophysiological mechanisms. In previous studies, we employed an integrated system of Transcranial Magnetic Stimulation and Electroencephalography (TMS-EEG) to track the effect of tDCS on cortical excitability. At rest, anodal tDCS (a-tDCS) over the right Posterior Parietal Cortex (rPPC) elicits a widespread increase in cortical excitability. In contrast, cathodal tDCS (c-tDCS) fails to modulate cortical excitability, being indistinguishable from sham stimulation.

Here we investigated whether an endogenous task-induced activation during stimulation might change this pattern, improving c-tDCS effectiveness in modulating cortical excitability.

In Study 1, we tested whether performance in a Visuospatial Working Memory Task (VWMT) and a modified Posner Cueing Task (mPCT), involving rPPC, could be modulated by c-tDCS. Thirty-eight participants were involved in a two-session experiment receiving either c-tDCS or sham during tasks execution. In Study 2, we recruited sixteen novel participants who performed the same paradigm but underwent TMS-EEG recordings pre- and 10 minutes post-sham and c-tDCS.

Behavioral results showed that c-tDCS significantly modulated mPCT performance compared to sham. At a neurophysiological level, c-tDCS significantly reduced cortical excitability in a frontoparietal network involved in task execution. Taken together, our results provide evidence of the state dependence of c-tDCS in modulating cortical excitability effectively. The conceptual and applicative implications are discussed.

## 1. Introduction

Over the past decades, tDCS has emerged as a non-invasive, cheap, safe, and easy-to-use technique to modulate cortical excitability in healthy volunteers and patients (Berryhill & Martin, 2018; Fregni et al., 2020). Despite its massive use, knowledge of tDCS physiological mechanisms remains incomplete.

Unlike Transcranial Magnetic Stimulation (TMS), which can induce post-synaptic excitatory potentials, tDCS is a neuromodulatory technique. Indeed, the current reaching the cortical surface is too weak to generate action potentials per se. Still, it is enough to alter the excitability and the spontaneous neural firing rate during and after the end of stimulation (Bocci, Ferrucci, & Priori, 2020; Nitsche et al., 2008; Stagg, Antal, & Nitsche, 2018). It is common knowledge that the modulatory effects of tDCS depend on the polarity of stimulation, with anodal tDCS (a-tDCS) increasing firing rate, hence excitability, and cathodal (c-tDCS) leading to the opposite outcome (Bindman, Lippold, & Redfearn, 1964; Creutzfeldt, Fromm, & Kapp, 1962; Nitsche & Paulus, 2000; Purpura & McMurtry, 1965). This physiological evidence has wrongly been extended to behavioral and psychophysiological measures, oversimplifying the cerebral dynamics underlying complex responses (Bradley, Nydam, Dux, & Mattingley, 2022). Evidence is indeed controversial concerning the established anodal-excitatory and cathodal-inhibitory effects (Jacobson, Koslowsky, & Lavidor, 2012; Schroeder & Plewnia, 2017). Studies investigating mechanisms underneath the complex pattern of reported effects are still unclear. Much literature neglected this issue and opted for using a-tDCS to explore brain-cognition interface or to treat different pathological conditions (Dedoncker, Brunoni, Baeken, & Vanderhasselt, 2016) since it results more effective in modulating behavior (Jacobson et al., 2012),

Among the advantages of using tDCS compared to TMS is the possibility of synchronously combining brain stimulation with a concurrent behavioral or cognitive task/training with minor exogenous distractions and somatosensory sensations. This option seems particularly convenient, considering that several studies highlighted that NiBS effects are *state-dependent;* namely, they are influenced by the current ongoing activity of the stimulated regions (Bikson & Rahman, 2013; Pisoni et al., 2018; Siebner, Hartwigsen, Kassuba, & Rothwell, 2009). Evidence from in vivo and in vitro studies indicates that a synaptic activity simultaneous with the stimulation is needed to induce detectable effects of tDCS (Cambiaghi et al., 2010; Priori, Berardelli, Rona, Accornero, & Manfredi, 1998). State-dependency mechanisms are largely unknown, although they could have crucial implications for basic research and clinical translation. According to the network activity-dependent hypothesis, tDCS effects would modulate neurons already engaged in an ongoing activity because they are close to the discharge threshold (Siebner et al., 2009). Therefore, stimulation delivered concurrently with a task recruiting the same neural network may induce a synergistic relationship between the endogenous neural activity generated by task execution and the exogenous source of activity promoted by the tDCS (Martin, Liu, Alonzo, Green, & Loo, 2014; Ohn et al., 2008; Stagg & Nitsche, 2011). Similarly, Bikson & Rahman (Bikson & Rahman, 2013) suggest that a functional property of tDCS is “activity-selectivity”, referring to the preferential modulation of the activated rather than inactive neuronal networks. Such property would also explain how tDCS can induce task-specific changes in brain activity despite being considered a technique with a low spatial focality. Critically, neurophysiological evidence in this sense has been provided in both the animal model and humans (Fritsch et al., 2010; Pisoni et al., 2018).

The concept of state dependency might represent a step forward in understanding many aspects of tDCS use. For instance, the influence of brain state in modulating tDCS effects can help explain polarity asymmetries and the variability of tDCS effects (Li, Uehara, & Hanakawa, 2015; Ridding & Ziemann, 2010; Vergallito, Feroldi, Pisoni, & Romero Lauro, 2022). Studies are indeed typically heterogeneous considering the coupling of brain stimulation and cognitive tasks, sometimes delivering tDCS before the task (as priming), sometimes during (as synergistic), and more rarely after (as consolidator) (Tatti et al., 2022). State dependency might also play a role in the reported lack of effectiveness of c-tDCS. Indeed, applying c-tDCS at rest (as in offline protocols) may fail to induce modulation in brain activity and behavior since a certain background activity might be needed to detect such an effect (Bortoletto, Pellicciari, Rodella, & Miniussi, 2015; Matsunaga, Nitsche, Tsuji, & Rothwell, 2004).

In a recent study combining tDCS with functional MRI (fMRI), Li et al. (Li et al., 2019) designed a factorial experiment manipulating the cognitive state (Choice Reaction Task vs. rest condition) and the stimulation polarity (anodal, cathodal, and sham). They delivered a short stimulation protocol over the right inferior frontal gyrus. When applied at rest, a-tDCS had a more pronounced effect compared to c-tDCS, increasing activation within the default mode network (DMN), which is typically active when individuals are not engaging in a specific activity or task, and decreasing activation of the salience network, which responds the subjective salience of stimuli (internal or external). C-tDCS effects were greater during task execution, where both stimulations increased SN activity.

The pattern highlighted by Li et al. partially aligns with previous studies from our research group. We assessed brain excitability and effective connectivity of cortical networks induced by tDCS using a system combining TMS with electroencephalography (TMS-EEG). Specifically, at resting state, a-tDCS over the right posterior parietal cortex (rPPC) elicited a widespread excitability increase along a bilateral frontoparietal network, likely overlapping the DMN (Romero Lauro et al., 2016, 2014). Conversely, despite maintaining the same parameters (duration, intensity, electrode size, and montage), c-tDCS failed to modulate cortical excitability, being indistinguishable from sham stimulation (Varoli et al., 2018). These results thus confirmed the reported imbalance of a-tDCS vs. c-tDCS (Dedoncker et al., 2016; Jacobson et al., 2012) at the resting state. Notably, when a-tDCS was delivered over the left inferior frontal gyrus during a fluency task, the increase in excitability rather than widespread was selective for the brain regions involved in task performance (Pisoni et al., 2018). Moreover, a-tDCS improved behavioral performance, and such enhancement positively correlated with changes in excitability.

In the present study, we aimed to complement this evidence by coupling c-tDCS over rPPC with concurrent tasks involving the stimulated brain network. To do so, we selected two visuospatial tasks: one tapping visual working memory (VWMT) (Heimrath, Sandmann, Becke, Müller, & Zaehle, 2012; Vogel & Machizawa, 2004) and the other involving attention reorienting, namely a modified version of the Posner Cueing task (mPCT) (Arif et al., 2020; Spooner, Wiesman, Proskovec, Heinrichs□Graham, & Wilson, 2020). Tasks were chosen because the rPPC seems to be a junction region between the cerebral circuits underlying visuospatial working memory and visuospatial orienting attention (Chica, Bartolomeo, & Lupiáñez, 2013; Juan, Tseng, & Hsu, 2017). During visual working memory tasks, brain activation has been shown to co-occur across the rPPC and the dorsolateral prefrontal cortex (Friedman & Goldman-Rakic, 1994). On the other side, neurophysiological, neuroimaging, and neuropsychological studies have consistently supported the critical role of rPPC in directing attention (Behrmann, Geng, & Shomstein, 2004; Corbetta, Kincade, Ollinger, McAvoy, & Shulman, 2000; Corbetta & Shulman, 2002; Thakral & Slotnick, 2009; Vallar & Perani, 1986).

We ran a first study in which cathodal and sham stimulation were delivered over rPPC during task execution to assess whether c-tDCS could induce a behavioral effect on participants’ performance. In a second experiment, we employed TMS-EEG in a new sample of participants to track the impact of c-tDCS combined with cognitive tasks on cortical excitability. We kept the same stimulation parameters to compare the results with our previous studies directly. We expected to detect, along with the behavioral modulation, a neurophysiological effect of c-tDCS, in contrast with the lack of effects observed when applied at the resting state.

## 2. Study 1

### 2.1 Materials and Methods

#### 2.1.1 Participants

Thirty-eight students (fourteen males, mean age = 22.8 years, SD = ± 4.1) participated in the study. Participants were right-handed according to the Edinburgh Handedness Inventory (Oldfield, 1971) (M = 0.88, SD = ± 0.13) and had no contraindications to tDCS (Rossi, Hallett, Rossini, Pascual-Leone, & Group, 2009). The experiment occurred in the Department of Psychology at the University of Milano-Bicocca. The Ethics Committee approved the study, and the participants’ treatment followed the principles stated in the Declaration of Helsinki.

#### 2.1.2 TDCS stimulation

TDCS was delivered by a BrainSTIM standard stimulator (EMS, Bologna, Italy). The intensity was 0.75 mA, and the duration was 15 min (30 s for sham), with 10 s of fade-in/fade-out period. The cathode (9 cm^2^, density .08 A/m^2^) was placed over the rPPC (P2), and the anode (25 cm^2^, density .03 A/m^2^) over the left supraorbital area.

#### 2.1.3 Tasks

##### 2.1.3.1 Modified Posner Cueing Task

In the mPCT, the trials followed the subsequent structure: i) a fixation point was positioned at the center of a black screen, flanked by two squares on the left and right sides; ii) after an interval ranging from 200-700 ms, the outline of one of the two squares became red (cue) for 100 ms; iii) 100 ms after the cue disappeared, a small gray square (target) appeared inside the left or right square and remained on the screen until participant response for a maximum time of 2000 ms (see Figure 1 – Panel A for a graphical representation). During the task, participants were instructed to focus on the fixation point and detect the target as accurately and fast as possible, pressing the letter “F” on the Italian keyboard for targets appearing in the left square and “J” for stimuli presented to the right.

**Figure 1.**
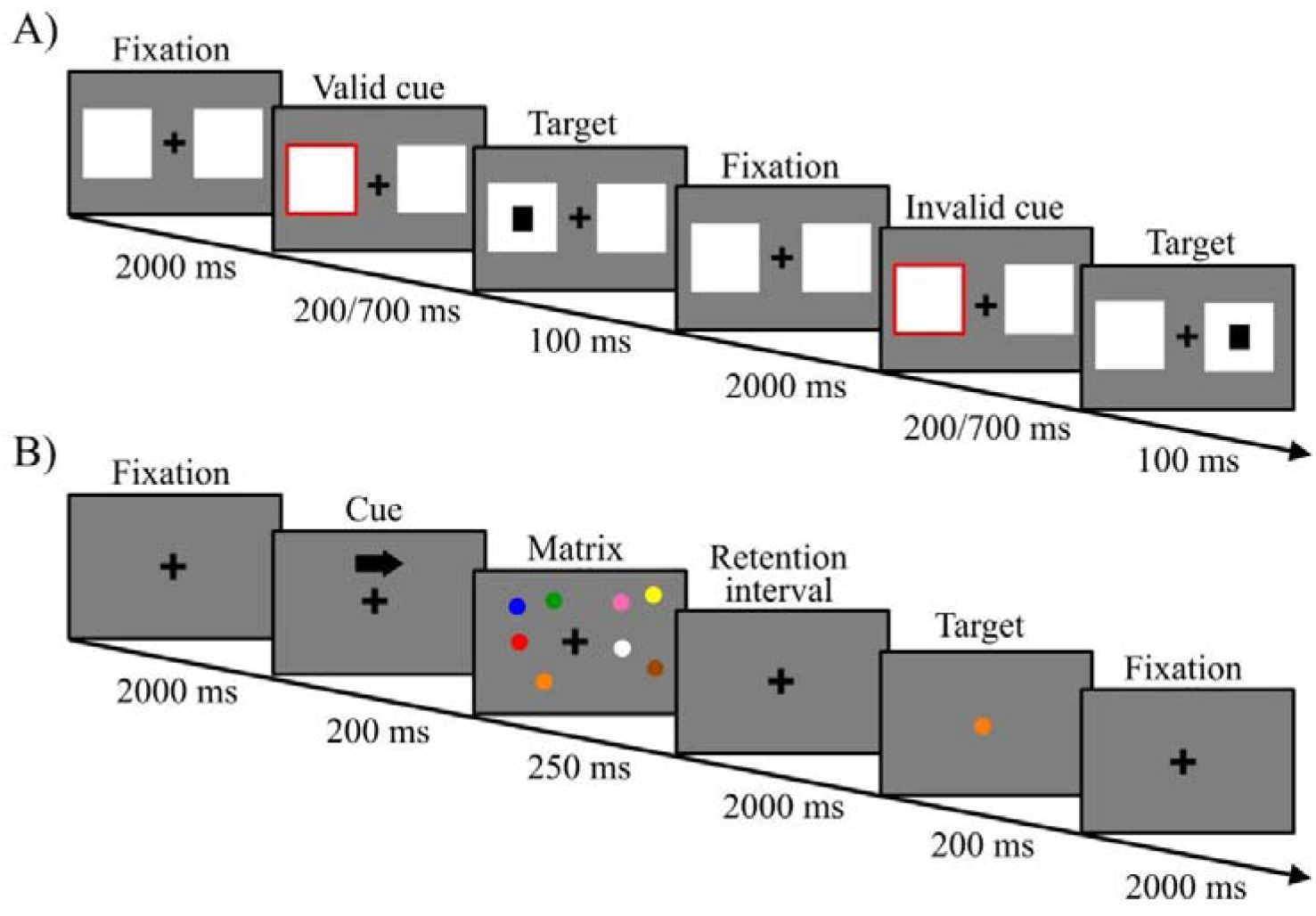
Graphical representation of the tasks’ procedure. Panel A highlights the mPCT procedure, with an example of valid (the first) and invalid (the second) trials. Panel B represents a trial of the VWMT in which the target was included in the previous matrix.

Two combinations of cue and target locations were possible: a valid or congruent condition, in which the target appeared on the same side of the bright cue, and an invalid or incongruent condition, in which the target and the cue appeared on the opposite sides. The mPCT comprised 48 congruent (24 on the right side and 24 on the left side of the central cross) and 48 incongruent trials (24 on the right and 24 on the left side of the central cross). Ten false alarm trials were added, in which no targets followed the cues. False alarms were not analyzed since participants did not have to press any key. We collected data using E-Prime 2 Software (Psychology Software Tools, Pittsburgh, PA). Response accuracy and reaction times (RTs) were recorded.

Notably, in the mPCT we changed the proportion of valid and invalid trials compared to the classic version of PCT (cPCT): whereas, in the cPCT the typical ratio of valid and invalid trials is 80% and 20%, respectively (Posner, 1980; Posner, Nissen, & Ogden, 1978), the mPCT comprised an equal amount of valid and invalid trials (Arif et al., 2020; Spooner, Wiesman, Proskovec, Heinrichs□Graham, & Wilson, 2020). The rationale for increasing the number of invalid trials in the mPCT, compared to the cPCT, was that we were planning to use the task in the subsequent TMS-EEG study. For methodological reasons, we then needed a sufficient and comparable number of trials for valid and invalid conditions to achieve reliable TMS Evoked Potentials (TEPs). We validated the mPCT in a pilot experiment described in the Supplementary materials (Section A).

##### 2.1.3.2 Visual Working Memory Task

The VWMT consisted of two blocks of 48 trials each, separated by a 30 s time interval. In each trial, participants maintained their gaze fixed on a central cross. After 200 ms, an arrow appeared on the top of the fixation point, and it could point either to the left or to the right hemifield, followed by a visual matrix comprising eight colored dots (size = 80 pixels; diameter = 1.75 cm), half localized on the right side and half on the left side of the screen. Within any visual matrix, all eight dots appeared with a different color and in a different position, varying within a rectangular portion of the hemifield. The colors and the positions were randomly assigned to avoid the compresence of two or more dots with the same shade and to reduce any facilitation effects. The visual matrix lasted 250 ms on the screen and was followed by a retention interval of 2000 ms. After that, a colored dot (target stimulus) appeared in concomitance with the fixation point, and participants had 2000 ms to establish if the displayed target was included (80% of the trials) or not (20%) in the initial matrix taking into consideration its color and ignoring its position (see Figure 1 – Panel B for a graphical representation). Participants were asked to press two keys on the keyboard to perform their choices using their index fingers. Due to the great difficulty of the task, evaluated by preliminary pilot subjects, participants completed a training session of 48 trials.

#### 2.1.4 Experimental procedure

Participants underwent two sessions, at least 48 hours apart (Nitsche et al., 2008), which differed only in the type of stimulation (c-tDCS or sham) they received. In each session, they performed the mPCT and the VWMT in a counterbalanced order. Both tasks were performed using a laptop (15.7” screen) at 60 cm from participants. The order of stimulation conditions was counterbalanced across participants.

#### 2.1.5 Statistical approach

MPCT and VWMT were analyzed separately. We performed analyses in the statistical programming environment R (R Core Team, 2022).

For the mPCT, we analyzed the dichotomous variable accuracy using general mixed effects models (Baayen, Davidson, & Bates, 2008), fitted using the GLMER function of the lme4 R package (Bates, Maechler, Bolker, & Walker, 2015). RTs values were analyzed using linear mixed-effects regression using the LMER procedure in the same lme4 package. The *trial validity* (factorial, two levels: valid vs. invalid), *tDCS condition* (factorial, two levels: cathodal vs. sham), and their interaction were entered in the full model as fixed factors. Moreover, we added the simple effect of the *trial order* to account for changes in performance due to learning or fatigue effects. A by-subject random intercept was included to account for participant-specific variability (Baayen et al., 2008). Fixed predictors’ inclusion in the final models has been tested with a series of likelihood ratio tests by progressively removing parameters that did not significantly increase the overall model goodness of fit (Gelman & Hill, 2006). For RTs, the automatic procedure step was used, while for the accuracy variable, this procedure was performed manually. Only RTs from accurate responses were included in the analyses, and outliers were removed using the model criticism.

Concerning the VWMT, in line with previous studies (Heimrath et al., 2012), we calculated the WM capacity K factor (Cowan, 2001; Pashler, 1988). This index assesses the individual WM performance for each *tDCS condition* (cathodal vs. sham) and the *attending hemifield* (left vs. right). K factor was estimated for each subject using the following formula: K = S (H – F). In the procedure, K represents the number of items that can be held in WM from an array of S objects. It assumes that the target item would have been stored in memory with respect to K/S of trials such that the performance will be correct on K/S on the change trials (= hit rate H). This procedure also considers the false alarm rate F to correct for guessing. In our statistical procedure, the continuous index K was analyzed using linear mixed models. We included participants’ intercept as a random structure, while *tDCS condition* (two levels: cathodal and sham) and *attended hemifield* (two levels: left vs. right) plus their interaction were added as fixed factors in the full model. As for the mPCT, we applied the automatic procedure step to remove the parameters that did not increase the model’s goodness of fit. For details on the model selection, see Supplementary Materials – Section B.

## 2.2 Results

### 2.2.1 mPCT

All false alarms and the data from two participants were excluded from analyses due to the low level of accuracy in one of the experiment sessions (one participant performed accurately in 52% of trials and the other 49%). Therefore, we ran the statistical analysis for the dependent variable accuracy on 6912 data points.

The best-fitting model included the simple effect of *trial validity* and *trial order* (χ^2^_(1)_ = 8.3, p = .004). Considering the simple effect of *trial validity*, accuracy was higher in the valid (M = 99.4, SD ± 7.8) compared to invalid (M = 97.8, SD ± 14.8) condition (p < .001). Concerning the effect of *trial order*, performance linearly increased during the experiment performance (p = .004).

Concerning RTs, we ran the analyses on 6814 data points. The best-fitting model included the simple effect of *trial order* and the interactions between the *tDCS condition* and *trial validity* (χ^2^_(1)_ = 11.2, p = .001). Considering the effect of *trial order*, RTs decreased during task performance, showing a learning effect (χ^2^_(1)_ = 41.1, p < .001). Relative to the interaction between *tDCS condition* and *trial validity* (χ^2^_(1)_ = 11.2, p = .001), post-hoc analysis showed that in the sham condition, a difference was traceable between valid and invalid presentations, with faster RTs in the valid condition (p < .001) while no differences were traceable for the cathodal stimulation (p = .940). Moreover, in the valid condition, RTs were slower in the cathodal stimulation compared to the sham condition (p = .021). Conversely, in the invalid condition, RTs were faster in the cathodal vs. sham condition (p = .021) (see Figure 2 for a graphical representation).

**Figure 2.**
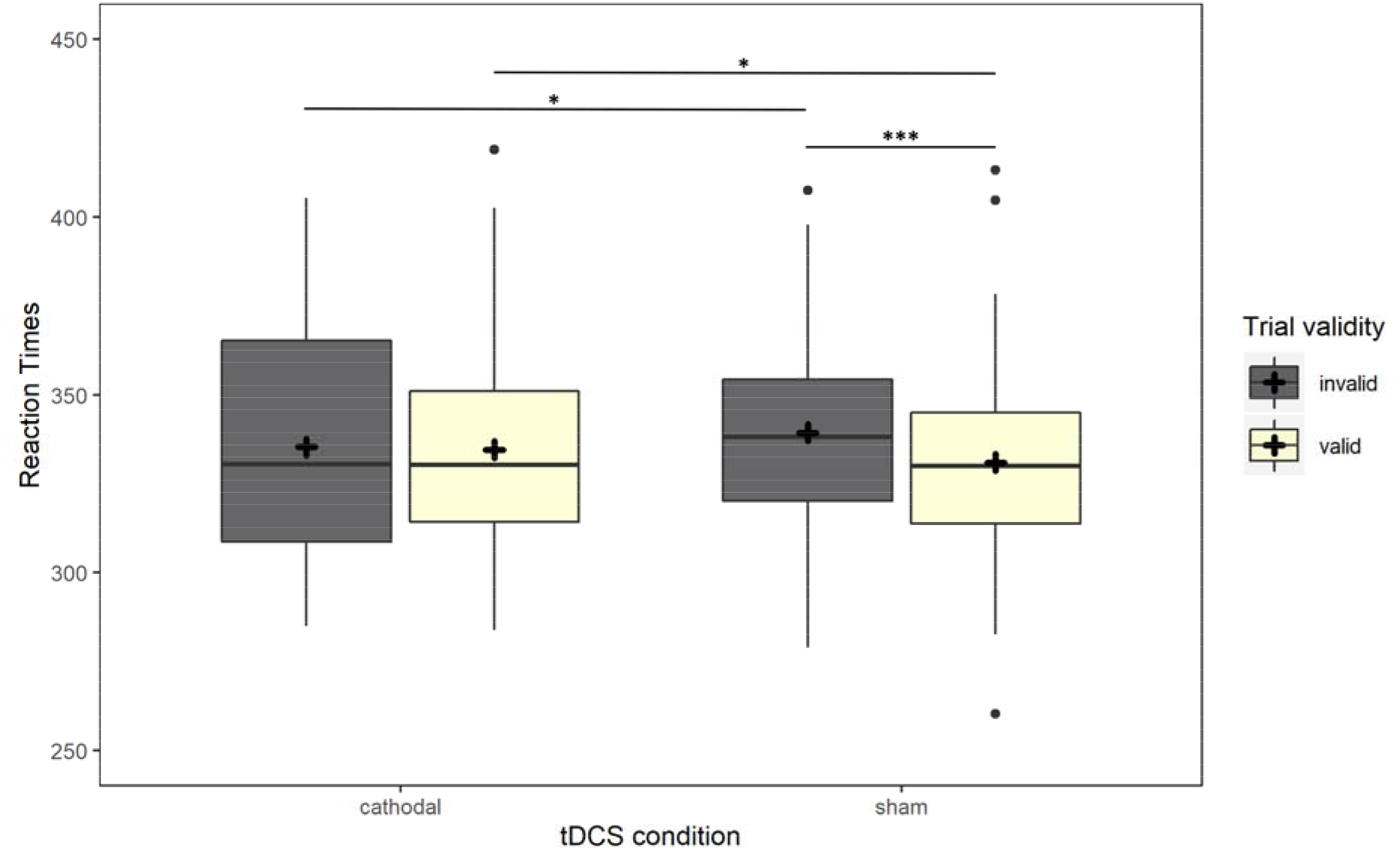
Results on RTs data in Experiment 1. The boxplots represent the RTs comparison between valid (light yellow boxes) and invalid (dark gray boxes) trial conditions in the c- and sham tDCS sessions, on the left and the right, respectively. Asterisks represent statistical p-values *** p < .001. ** p < .01, * p < .05.

#### 2.2.2 VWMT

One participant was removed due to a technical problem during task execution. The analysis was run on 148 data points. The best model was the one without a fixed factor (See Table S5). Therefore, no significant differences were found between the two stimulation conditions, the attended hemifields, nor their interaction.

### 2.3 Interim conclusions

The results from Study1 shed light on the feasibility of using c-tDCS over rPPC to modulate task performance in one of the two selected visuospatial tasks, namely mPCT. We observed indeed that c-tDCS abolished the advantage for valid cues. Once tested the behavioral effectiveness of c-TDCS over rPPC in modulating the performance at a visuospatial task, we could proceed to the Study 2 to explore the neurophysiological counterpart of these behavioral effects and to test our main hypothesis, i.e. whether neurophysiological effects on cortical excitability might arise when c-TDCS over rPPC in delivered during a task which involve this brain region to be executed.

## 3. Experiment 2: TMS-EEG

### 3.1 Materials and Methods

#### 3.1.1 Participants

Sixteen healthy volunteers (seven males, mean age = 25.4 years, SD = ± 3.5), different from those involved in Experiment 1, participated in Experiment 2. They were right-handed (mean laterality coefficient = 0.89, SD = ± 0.14) and were included following the same criteria as Experiment 1.

#### 3.1.2 Procedure

All volunteers participated in two experimental sessions, performed at least one week apart, corresponding to c-tDCS and sham conditions. The order of the tDCS conditions was counterbalanced across participants. Each session consisted of two blocks of TMS-EEG recordings performed before (pre-tDCS) and 10 min after the end of tDCS stimulation (post-tDCS) (see Figure 3 for a schematic representation of the procedure). Considering that c-tDCS affected performance only in the mPCT and that previous evidence suggested that tDCS has a cumulative effect on performance (Boggio et al., 2008; Monte-Silva, Liebetanz, Grundey, Paulus, & Nitsche, 2010), we decided to present the two tasks in a fixed order to maximize behavioral modulation. Therefore, in Experiment 2, participants performed first the VMWT and then the mPCT.

**Figure 3.**
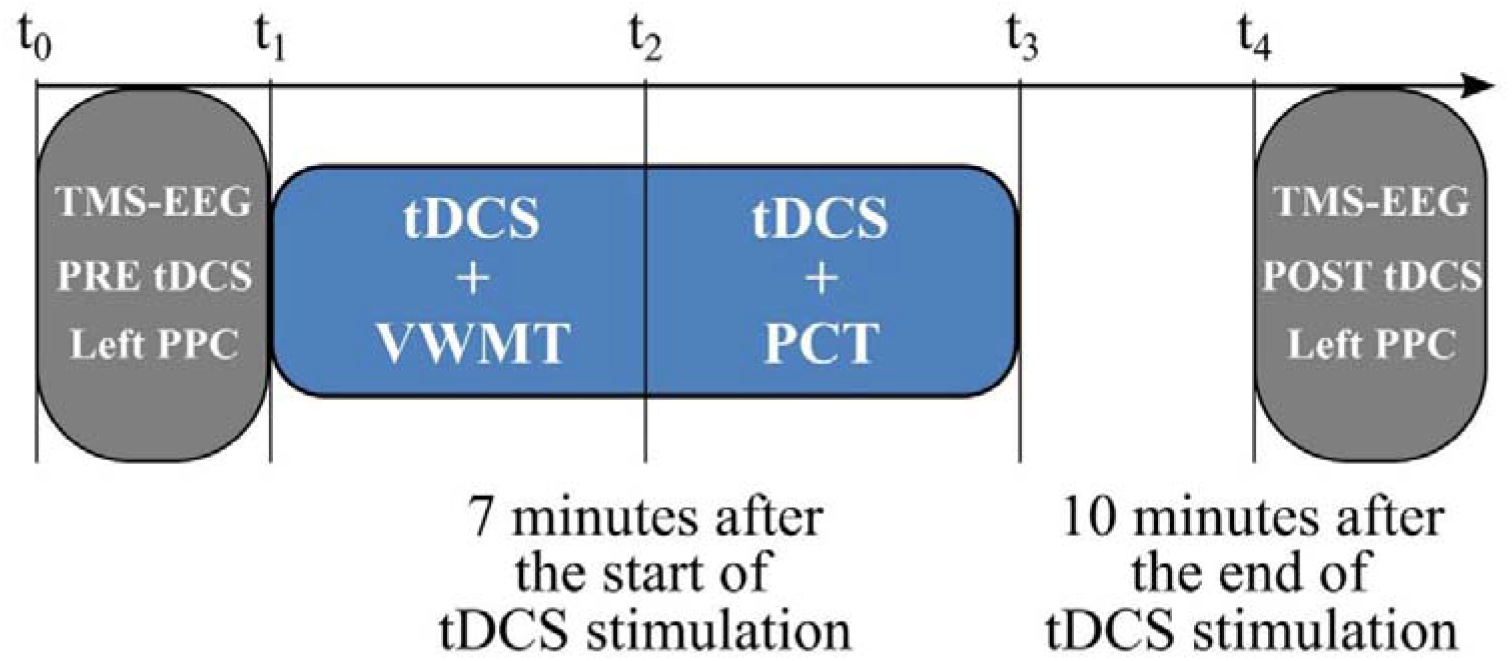
Graphical explanation of TMS-EEG experimental procedure.

#### 3.1.3 TDCS parameters

We applied the same stimulation parameters used in Experiment 1. The cathode was placed under the P2 EEG electrode, previously removed from the cap (see Figure 4 – Panel b) for the tDCS estimated electric field).

**Figure 4.**
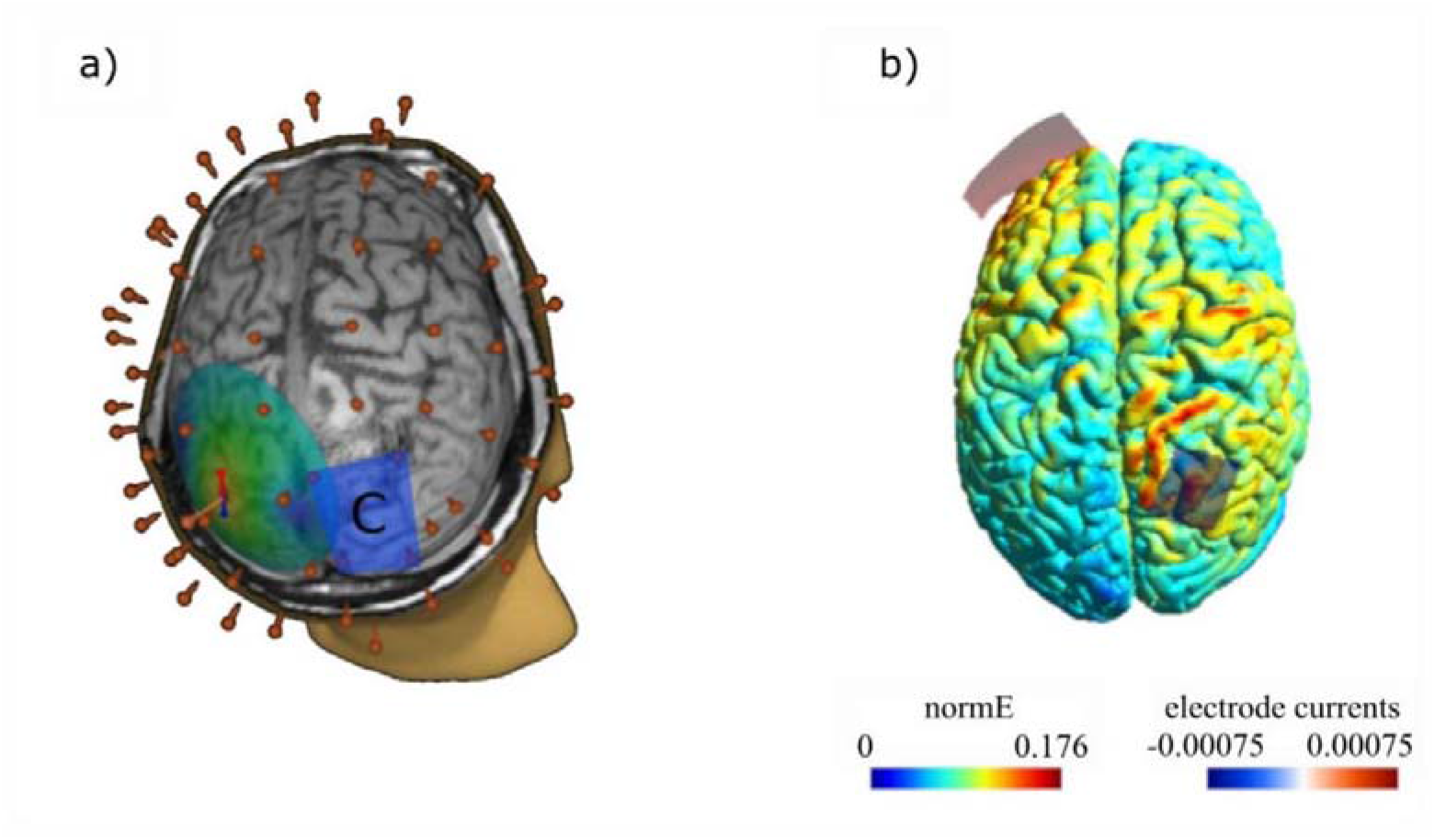
TDCS and TMS-EEG setting. Panel a: individual MRI 3D reconstruction showing the TMS electrical field on the left PPC (where TMS single pulses were delivered). The blue rectangle represents the cathode position over the rPPC; the 60 red points correspond to the EEG electrodes. Panel b: the tDCS estimated current flow.

#### 3.1.4 TMS-EEG parameters

For TMS-EEG recording, single pulses of TMS were delivered with an Eximia™ TMS stimulator (Nexstim™, Helsinki, Finland) using a focal figure-of-eight bi-pulse 70 mm-coil while concomitantly recording EEG from a 60 channels cap. The stimulation target was the left PPC, between P1 and CP1 EEG electrodes.

Coil positioning and monitoring throughout recordings were achieved using a Navigated Brain Stimulation (NBS) system (Nexstim™, Helsinki, Finland) on the individual high-resolution (1 × 1 × 1 mm) structural MRI. The MRIs were previously acquired for each participant using a 3T Intera Philips body scanner (Philips Medical Systems, Best, NL). The coil was placed tangentially to the scalp. The position was adjusted for each participant to direct the electric field perpendicularly to the shape of the cortical gyrus, following the same procedure used in previous studies (Casarotto et al., 2010; Mattavelli, Rosanova, Casali, Papagno, & Lauro, 2013; Romero Lauro et al., 2014) (see Figure 4 – Panel a for a graphical representation).

The average stimulation intensity, expressed as a percentage of the maximal output of the stimulator, was 64% (range = 60 - 73%), corresponding to an electric field of 105 ± 12 V/m. TMS-evoked potentials (TEPs) were recorded using single TMS pulses on the left PPC. Each recording session lasted about 7 min, during which 180 pulses were delivered, with an inter-stimulus interval (ISI) randomly jittered between 2,000 and 2,300 ms (0.4 – 0.5 Hz). During TMS-EEG registration, participants fixated on a white cross on a black screen (17”). Participants heard a noise-masking trace during the TMS-EEG recording, to avoid the presence of auditory artefacts (Casarotto et al., 2010).

#### 3.1.5 TMS-EEG data preprocessing

EEG data preprocessing was performed with Matlab R2016b (Mathworks, Natick, MA, USA). First, recordings were down sampled to 725 Hz. The continuous signal was split into single trials, from 800 ms before to 800 ms after the TMS pulse. Trials with artifacts due to eye blinks/movements or spontaneous muscle activity were removed following a semi-automatic procedure (Casali et al., 2010) and the visual inspection of the signal by trained experimenters. TEPs were computed by averaging selected artifact-free single trials and filtering them between 2 and 40 Hz. Bad or missing channels, such as P2 and CP2 that were right above the cathode were interpolated using the spherical interpolation function of EEGLAB (Delorme & Makeig, 2004) TEPs were then average-referenced, and baseline corrected between −300 and −50 ms before the TMS pulse. The average number of trials considered in the analysis was 122 (SD = ± 4) for the pre-tDCS and 120 (SD = ± 2) for the post-tDCS condition.

#### 3.1.6 TMS-EEG data analyses

We performed three analyses on our EEG dataset for c-tDCS and sham conditions. Two analyses were performed at the sensors level: Global and Local Mean Field Power (GMFP, LMFP) computation and cluster analysis. GMFP and LMFP were computed as index of global and local excitability and to mirror the analyses run in our previous studies in which a- and c-tDCS were delivered at resting state (Romero Lauro 2014; 2016; Varoli et al., 2018). We added the cluster analysis to avoid the selection of a priori time windows and regions of interest. The third analysis was performed at the level of cortical sources, to avoid the potential confound of volume conduction, even more relevant with the delivery of tDCS (Bailey et al., 2016) and to achieve a better definition of the spatial distribution of the tDCS effects (Casarotto et al., 2010; Romero Lauro et al., 2016). For the sake of brevity here we report the procedures and results of the Cluster analysis and source analysis. Details about the procedures and the results of GMFP and LMFP can be found in Supplementary Materials (Section C).

##### 3.1.6.1 Cluster analysis

For both cathodal and sham conditions, the pre-and post-tDCS sessions were compared through a cluster-based permutation test (Maris & Oostenveld, 2007) implemented in the “FieldTrip” MATLAB toolbox for M/EEG analysis (Oostenveld, Fries, Maris, & Schoffelen, 2011). This procedure solves the multiple comparisons problem by permuting and clustering data based on temporal and spatial proximity. More precisely, a big number N of data permutations are performed by shuffling the labels of the experimental conditions. Then t-tests are computed at each time point for each permutation. All samples with a statistic corresponding to a p-value smaller than .05 are thus clustered based on spatial proximity. Cluster-level statistics are calculated by taking the sum of the t-values within each cluster. Finally, the cluster-corrected threshold is computed as the maximum cluster-level statistics permutation distribution (Maris & Oostenveld, 2007). In these analyses, 10000 permutations were performed for each comparison with a permutation-significance level of p = 0.05 for the time window between 0 and 300 ms from the TMS onset. We chose this interval after observing the presence of relevant peaks in the butterfly plots, which occurred, as usual, in the first 300 ms (Casarotto et al., 2010).

##### 3.1.6.2 Source analysis

As the first step, individual standardized meshes were reconstructed for each participant starting from their structural MRIs (SPM8) (Ashburner, Barnes, & Chen, 2011). We obtained meshes of cortex, skull, and scalp compartments (containing 3004, 2000, and 2000 vertices, respectively), normalized to the MNI atlas. Then, the EEG sensor’s position was aligned to the canonical anatomical markers (pre-auricular points and nasion) for each participant, and the forward model was computed. The inverse solution was calculated on the average of all artifact-free TMS-EEG trials, using the weighted minimum norm estimate with a smoothness prior, following the same procedures as in Casali and colleagues (Casali, Casarotto, Rosanova, Mariotti, & Massimini, 2010). After source reconstruction, a statistical threshold was computed to assess when and where the post-TMS cortical response differed from pre-TMS activity. We applied a non-parametric permutation-based procedure (Pantazis, Nichols, Baillet, & Leahy, 2003). We obtained a binary spatial-temporal distribution of statistically significant sources, including only information from significant cortical sources in further analyses. As a measure of global cortical activation, we cumulated the absolute significant current density (global SCD, measured in μA/mm^2^) (Casali et al., 2010) (overall 3,004 cortical vertexes) for each recording session (pre-tDCS, post-tDCS). To mirror the LMFP at the sensors level, we computed a local SCD in the vertexes within four different Brodmann’s areas (BAs), identified using an automatic tool of anatomical classification (WFUPickAtlas tool, http://www.ansir.wfubmc.edu). These BAs corresponded to the original four clusters of LMFP (left/right BA6 and 7).

To coherently perform the analysis at the level of the sources, we started with the sensor cluster analysis results. The global SCD and local SCD were then computed in the time window from 180 to 230 ms, in which the significant cluster was found. On this data, a linear mixed model with the factor *condition* (Pre-tDCS vs. post-tDCS) was run for Global SCD and the four ROIs (left and right BA7 and BA6). We used the same analyses and procedures to analyze data from sham sessions.

### 3.2 Results

#### 3.2.1 Behavioral results

Behavioral data were analyzed following the same procedure described in Experiment 1. Details of model selection are reported in the Supplementary materials (Section C).

MPCT statistical analyses were performed on 3072 data points. The best-fitting model for the dependent variable accuracy included the simple effects of *trial order* and *trial validity* (χ^2^_(1)_ = 5.6, p = .018) (see Table S6 for the model selection). Considering the *trial order* effect (χ^2^_(1)_ = 5.7, p = .017), participants became progressively less accurate in the mPCT performance, suggesting an effect of fatigue during task execution. Concerning *trial validity* (χ^2^_(1)_ = 13.4, p < .001), as expected participants’ accuracy was higher in the valid vs. invalid condition. Relatively to RTs, we considered only accurate responses, and analyses included 3014 data points. The best-fitting model comprised the simple effects of *trial order* and *tDCS condition* (χ^2^_(1)_ = 34.4, p = < .001) (see Table S7 for details on model selection). Considering the effect of *trial order* (χ2(1) = 34.6, p < .001), participants became faster while performing the task, showing an effect of learning on the performance speed. Relatively to the *tDCS condition* (χ^2^_(1)_ = 9.6, p = .002), RTs were slower during the cathodal as compared to the sham stimulation (see Figure 5 for a graphical representation). Considering the VWMT, statistical analyses were performed on 64 data points. The results showed no significant main effects or interaction between the *tDCS condition* and *attended hemifield* (see Table S8 for details on model selection).

**Figure 5.**
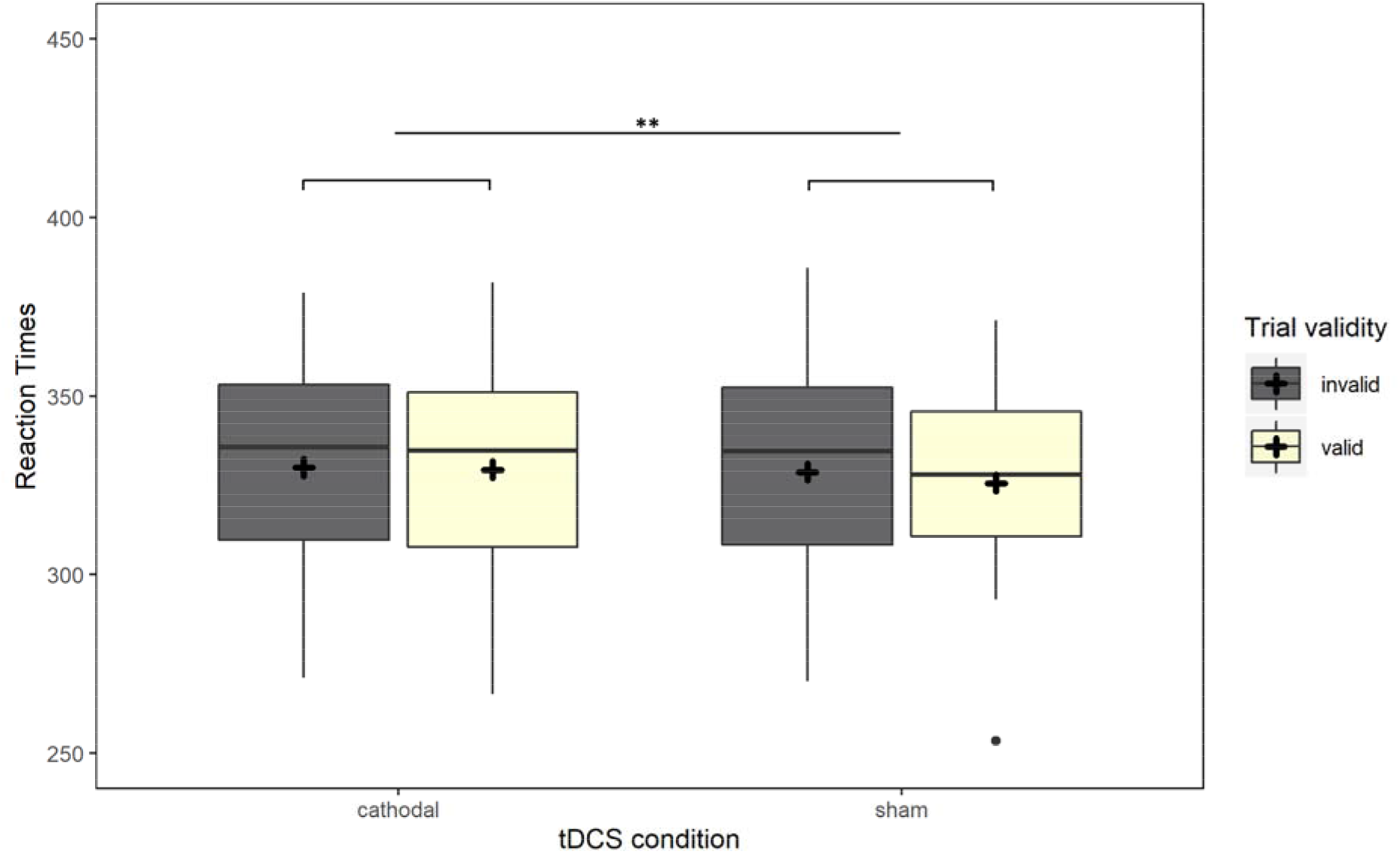
Behavioral results of mPCT in Experiment 2. The boxplots represent the RTs data for the valid (light yellow boxes) and invalid (dark gray boxes) trial conditions in the c- and sham tDCS sessions, on the left and the right, respectively. Asterisks represent statistical p-values *** p < .001. ** p < .01, * p < .05.

#### 3.2.2 TMS-EEG data

We here report the results of cluster and source analysis. Results of GMPF and LMFP can be found in the Supplementary materials – Section C.

##### 3.2.2.1 Cathodal Stimulation - Cluster analysis

A cluster-based analysis evaluating the effect of tDCS stimulation (Pre-tDCS vs. Post-tDCS) revealed a significant positive cluster (p < .023; Pre > Post) in frontoparietal electrodes in a time range from 180 to 230 ms from the TMS onset. TEPs’ scalp topographies of statistically significant differences showed that the positive cluster was associated with frontocentral electrodes and covered a bilateral portion of brain regions.

##### 3.2.2.2 Cathodal Stimulation - Source modeling analyses

Source modeling analyses performed in the cluster analysis’ significant time window confirmed results from the cluster analysis at the sensors level. The best-fitted model on the index of global cortical activation, namely Global SCD, included the main effect *condition* (χ^2^_(2)_ = 6.4; p = .014) with a decrease of TEPs amplitude after tDCS stimulation (see Figure 7). Similarly, in the analysis for the Local SCD computed for 4 BAs, namely bilateral BA6 and BA7, the factor *condition* was included in the final model as the main effect for left BA7 (χ^2^_(2)_ = 5.87; p = .015), right BA7 (χ^2^_(2)_ = 6.0; p = .014) and left BA6 (χ^2^_(2)_ = 7.9; p = .005). Only for the right BA6 no effect of real c-tDCS was highlighted on local SCD (χ^2^_(2)_ = 1.5; p = .224).

##### 3.2.2.3 Sham stimulation - Cluster analysis

For the sham stimulation condition, the results revealed neither positive nor negative significant clusters (Figure 6).

**Figure 6.**
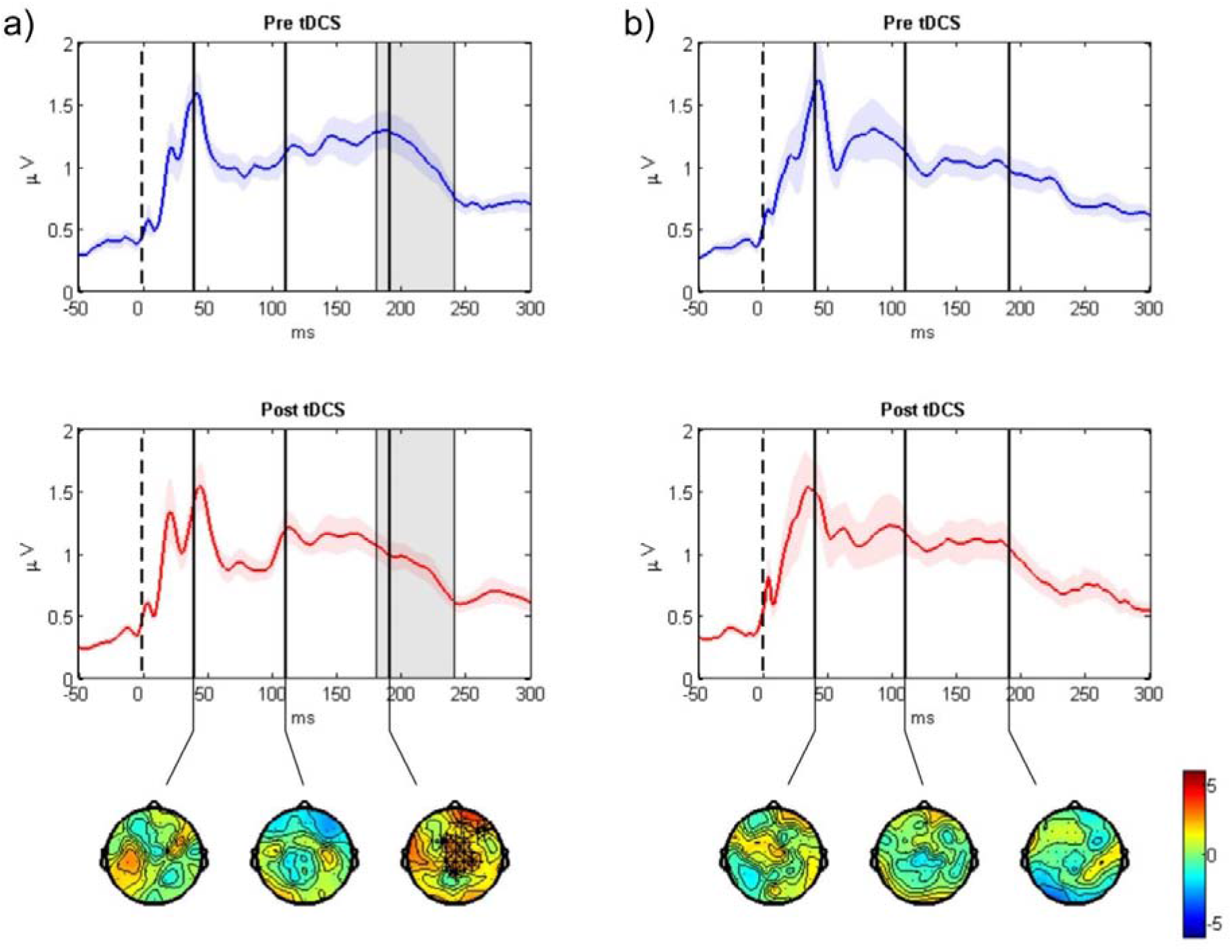
Results of cluster analysis. GMFP plot and topoplot of the Pre-tDCS (blue trace) and Post-tDCS (red trace) sessions for c- and sham tDCS, on panels a and b, respectively. The gray bars define the time window where cluster analysis evidenced significant results. In the last row, topoplots show the significant clusters evidenced by the cluster analysis.

##### 3.2.2.4 Sham Stimulation - Source modeling analyses

We decided to perform the Source modeling analysis to explore at a different signal level, although no significant results were found at the sensors level with cluster permutation. For this reason, the Source modeling analyses were performed between 180 and 230 ms, which is the significant time window for c-tDCS. The sham data lacked statistically significant effects, confirming the results of the sensor analyses, but at a global level only. The Global SCD final model did not include *condition* (χ^2^_(2)_ = 0.13; p = .711), indicating no effect of sham stimulation on cortical excitability (see Figure 7).

**Figure 7.**
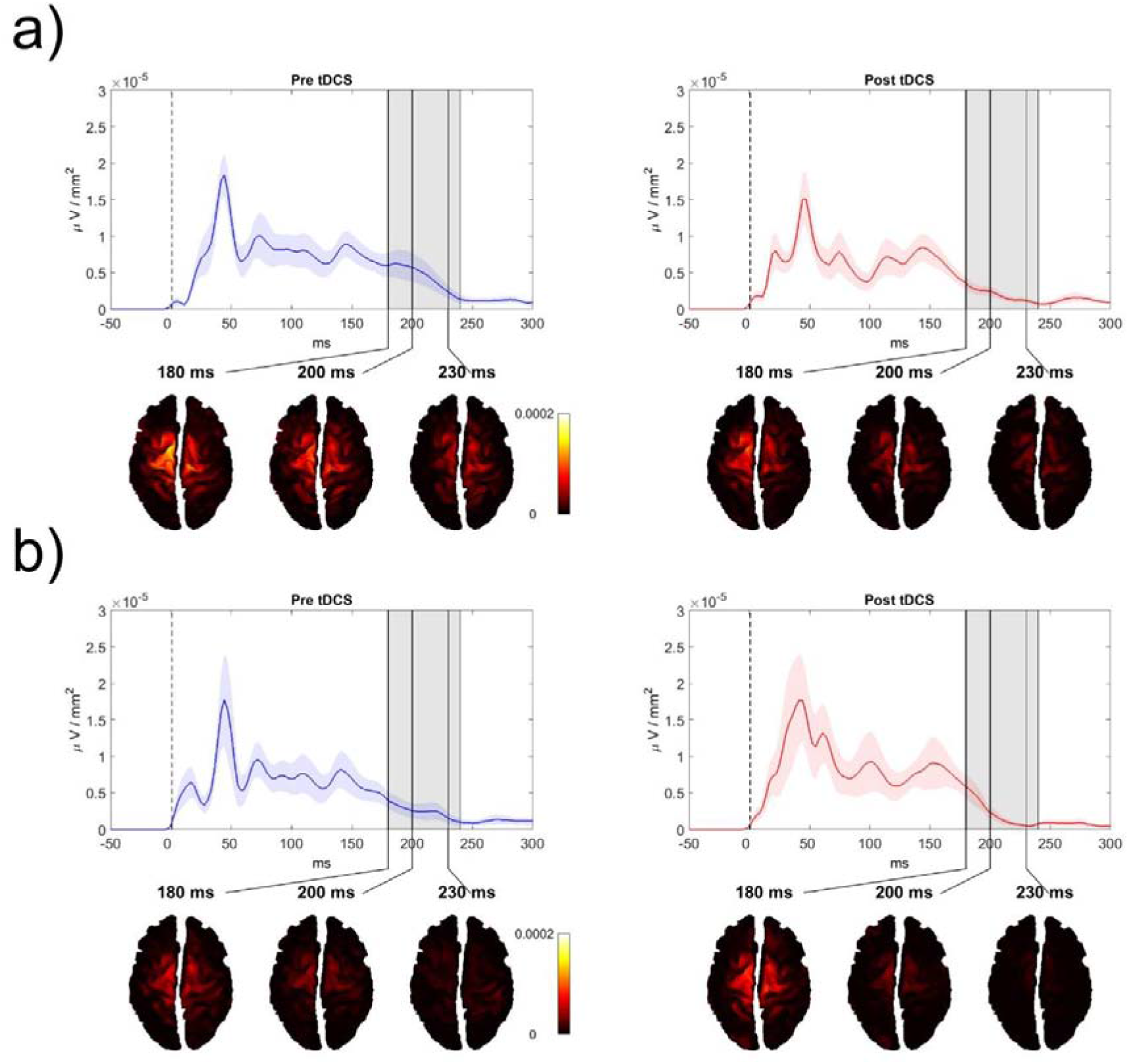
Active vertexes and current spread at the local maxima in the TEPs time windows were revealed by the cluster analysis for both c- and sham tDCS on panels a and b, respectively. For each recording session (pre-and post-tDCS), the GMFP is shown on the first top row. The gray bars define the time window where cluster analysis evidenced significant results. The second row shows the estimated cortical sources in time coincidence with the maximum GMFP value, between 180 and 230 ms.

The same holds for both left (χ^2^_(2)_ = 0.03; p = .850) and right (χ^2^_(2)_ = 0.15; p = .708) BA7, as well as for left BA6 (χ^2^_(2)_ = 0.56; p = .450), where LRT indicated no inclusion of the factor *condition* in the final model. Different from what emerged for sensors data, at the level of the sources, the main effect of the *condition* was significant (χ^2^_(2)_ = 5.19; p = .022) for the right BA6. Results in this area showed wider TEPs after sham stimulation than those before.

Finally, as in a previous study by Pisoni et al. (Pisoni et al., 2018), we investigated possible links between cognitive and neurophysiological measures by computing correlations between the real vs. sham differences in the mPCT performance (RTs) and cortical excitability. Results did not highlight significant correlations between changes in excitability and behavioral performance (ps > .204) (see supplementary materials Figure S4 for a graphical representation of the correlations pattern).

### 3.3 Interim Comment

Regarding the behavioral results, we confirmed that c-TDCS over rPPC affected the performance at mPCT and not VWMT. We partially replicated the behavioral effects in mPCT observed in Study 1, since in Study 2 c-TDCS slowed down the RTs compared to sham stimulation, independently from the trial validity. Crucially, at a neurophysiological level, cluster analysis and source analysis converged in showing that c-tDCS reduced cortical excitability in the time window of 180-230 ms from TMS onset, in a frontoparietal network.

## 4. General Discussion

The present study investigated the behavioral and neurophysiological effects of c-tDCS delivered over rPPC during the performance of two visuospatial tasks.

Concerning the behavioral effect, the results of Experiment 1 and 2 converge in showing that c-tDCS on rPPC reduced proficiency in the mPCT. Critically, we observed a significant increment of RTs in the mPCT during c-tDCS compared to the sham condition. More specifically, c-tDCS eliminated the advantage for RTs in the valid cue condition, since no difference was found between valid and invalid cues, as observed in the sham condition and the pilot study. These results suggest that tDCS acted in reducing the pre-allocation of attention elicited by an exogenous cue. Previous studies investigating tDCS effects on visuospatial attention showed mixed results, thus providing inconclusive findings (Demartini et al., 2019; L. M. Li, Leech, et al., 2015; Lo, van Donkelaar, & Chou, 2019; Roy, Sparing, Fink, & Hesse, 2015). It is possible that the methodological differences, such as montage, polarity, and timing of stimulation, accounted for this heterogeneity in results. C-tDCS did not impact performance on the VWMT, neither in Experiment 1 nor 2. Also in this case, previous results were mixed, with some studies highlighting a modulatory tDCS effect in this domain, and others not (Heimrath et al., 2012; Hill, Fitzgerald, & Hoy, 2016; Robison, McGuirk, & Unsworth, 2017; Tseng et al., 2012). Results inconsistency has been previously related to the strategies employed in the task execution, which largely depend on the individual WM capacity (Heinen et al., 2016; Jones & Berryhill, 2012; S. Li et al., 2017).

Concerning the neurophysiological data, the cluster analysis at the sensor level revealed a significantly different bilateral frontoparietal positivity comparing pre- and post-c-tDCS, between 180 and 230 ms from the TMS onset: cortical excitability decreased after c-tDCS compared to the pre-stimulation recording. Source modeling analysis confirmed the cluster analysis results. Indeed, SCD changed after stimulation compared to the pre-tDCS condition when it was computed at a global level and in three of the four BA. Computing the data in the time window derived from the cluster analysis (180 – 230 ms), a significant change was found bilaterally for BA7 and left BA6, but not for right BA6.

In a previous study (Varoli et al., 2018), we failed to observe any changes in cortical excitability when c-tDCS was delivered at rest. Since in this previous study the analyses were performed only in the first 150 ms of TEPs, to rule out that the absence of neurophysiological effects might depend upon a too short time window of interest, the same cluster analysis performed in the present study was also run on the previously collected dataset. Crucially no significant positive or negative clusters were found in the re-analysis.

Together with our previous studies, the present findings confirm the state dependency of tDCS effects. Crucially, here we provide clear-cut evidence that such state dependence is even more crucial in the case of cathodal stimulation to effectively modulate human cortical excitability outside the corticospinal excitability domain.. It can be argued, indeed, that c-tDCS - which should reduce the level of neural discharge - may produce effects only in systems with a high level of basal activity, as that elicited by concurrent task execution. It is possible that c-tDCS may have little or no effect at resting state because only low levels of spontaneous discharge occur. A possible explanation could be the crucial role played in tDCS effects by the location and frequency of active synapses, as suggested by in vitro modeling (Lafon, Rahman, Bikson, & Parra, 2017). The state-dependent effectiveness for neurophysiological modulation that we observed might be strictly connected to the long-lasting plastic changes induced in the stimulated area by the concurrent task execution.

The crucial role of state-dependency for c-tDCS neurophysiological effects might provide novel insights in the mechanism underlying anodal vs cathodal imbalance. Despite the increasing evidence on the state-dependency of NiBS effects, there is still inconsistency in literature about the application of tDCS during a task. Many scholars prefer to use tDCS as priming – namely at rest and before performing a task. The often-reported null effects of cathodal-tDCS might then depend on the frequency of offline paradigms. Jacbson et al (2012) systematically explored the factors underlying anodal vs cathodal imbalance of effects and proposed that a relevant role is played by the motor vs cognitive targeted function, being the likelihood of achieving anodal-excitatory vs cathodal-inhibithory effects higher in the former and reduced in the second (they also clarified that in the cognitive domain usually there is significant higher chance to observe an excitatory effect of anodal whereas a lower probability to observe an inhibitory effect of c-tDCS, usually leading to null results). Interestingly, they also checked whether several stimulation parameters, likewise the intensity, duration and electrodes size, might affect such imbalance, without finding any significant result. However, they did not take into consideration the role of online vs offline paradigm, i.e. whether the stimulation was applied at rest or before vs during a task. Our data suggest that status of activation of the stimulated brain area might affect such imbalance, and the prevision would be that a lower imbalance can be found in online paradigm. Future studies might further test this hypothesis.

Notably, the neurophysiological c-tDCS effects seem to be restricted to those areas activated by the task execution, as previously observed for a-tDCS (Pisoni et al., 2018). Different studies have indeed shown that a dorsal frontoparietal network is globally associated with attention orientation, even if the regions in the system are engaged differently over time and across the hemispheres concerning the type of attention (Chica et al., 2013). Within this framework, attentional performance critically depends on the interaction between PPC and the frontal eye fields, one holding the sensory representation, the other holding a motor representation (Mesulam, 1981; Torriero et al., 2019). In line with our results, Thiel et al. (Thiel, Zilles, & Fink, 2004) showed that the specific reorienting attention activity in PCT increased activation in a bilateral frontoparietal network, including left and right intraparietal sulcus, right temporoparietal junction, and left and right middle frontal gyrus in healthy volunteers. Our study found a decreased cortical excitability in the left but not in the right frontal area. Although frontal and parietal lesions can both induce neglect (Halligan, Fink, Marshall, & Vallar, 2003), there is some divergence in imaging and patient data concerning the role of the frontal cortex in reorienting attention since some frontal patients did not show specific deficits in these types of tasks (Petersen, Robinson, & Currie, 1989; Posner, Inhoff, Friedrich, & Cohen, 1987). The occurrence of neurophysiological effects in task-relevant brain areas support the activity-selectivity hypothesis, according to which functional tDCS specificity may derive from either active neuronal networks that are preferentially modulated by tDCS or from input-selectivity, where a bias is applied to different synaptic inputs (Bikson & Rahman, 2013).

Regarding the timing at which the neurophysiological effects of the stimulation were observed, c-tDCS affected cortical excitability at a relatively later stage. Whereas early TEPs are considered a direct and reliable marker of cortical excitability of the targeted area (Ilmoniemi & Kičić, 2010; Pellicciari, Brignani, & Miniussi, 2013), middle-latency component reflects the connectivity of functional network activated by the task (Casarotto et al., 2010; Casula et al., 2022; Ilmoniemi & Kičić, 2010; Pisoni et al., 2018; Veniero, Brignani, Thut, & Miniussi, 2011). The timing of neurophysiological effects might then suggest that rather than local excitability, c-tDCS affected the connectivity within the frontoparietal network underlying the attentional process required by mPCT.

Crucially, no significant changes in cortical excitability were found when sham stimulation was delivered. The only exception is at the level of the sources where, only in correspondence with the right BA6, we found an increase in SCD in the post-compared to pre-tDCS condition. What occurs in the sham control condition can be considered a sort of baseline, reflecting modulations possibly due to the task execution. Therefore, the significant increase of SCD in the right BA6 after sham may be attributed to the involvement of this area in the visuospatial orienting attention (Brovelli, Lachaux, Kahane, & Boussaoud, 2005; Nobre, Gitelman, Dias, & Mesulam, 2000). Consequently, the lack of effect on right BA6 after c-tDCS can be interpreted as an effect of the stimulation: c-tDCS might have reduced cortical excitability in this area, canceling the activation effect due to the task execution observed in the sham condition. It could be possible that the right BA6 plays a peculiar role in the excitability-inhibitory processes in the network. This area could compensate for the destructive effects of c-tDCS on the other regions in the system during task execution. Further studies need to be performed to disentangle this issue.

To conclude, our results shed light on the dependence of tDCS effects on the state of activity of the stimulated region. Supporting previous evidence for a-tDCS (Pisoni et al., 2018), the present findings showed that delivering c-tDCS during a task would confine the neurophysiological effect on the task-relevant areas, thus accounting for the specificity of the behavioral impact usually observed. In the case of c-tDCS, at least for parietal cortex stimulation, the concurrent task activity seems essential to observe cortical excitability modulations. The possibilities of translating these knowledge into a clinical domain are promising, considering that neurological and psychiatric conditions are characterized by the presence of a pathologically altered neural plasticity and connectivity, that could be restored by combining brain stimulation with concurrent cognitive tasks, training or psychotherapy (Dedoncker, Baeken, De Raedt, & Vanderhasselt, 2021; Sathappan, Luber, & Lisanby, 2019; Tatti, Rossi, Innocenti, Rossi, & Santarnecchi, 2016; Vergallito et al., 2021).

## Open practices statement

The datasets generated and the script for the analyses are publicly available at https://osf.io/7jdmp/. The experiment was not preregistered.

## Notes

https://osf.io/7jdmp/

### Competing Interest Statement

The authors have declared no competing interest.

### Summary of Updates

Figure S1 supplementary materials changed

