## Supplementary material for "State-dependent effectiveness of cathodal transcranial direct current stimulation on cortical excitability": Supplementary_Materials_0603.pdf

#### 1. Section A – Pilot experiment

The pilot experiment was implemented to compare participants' performance in a modified (mPCT) vs. classic version of the Posner Cueing task (cPCT). In the mPCT, we changed the proportion of valid and invalid trials compared to the cPCT: whereas, in the cPCT the typical ratio of valid and invalid trials is 80% and 20%, respectively (Posner, 1980; Posner, Nissen, & Ogden, 1978), the mPCT comprised an equal amount of valid and invalid trials (Arif et al., 2020; Spooner, Wiesman, Proskovec, Heinrichs-Graham, & Wilson, 2020).

##### 1.1. Materials and methods

###### *1.1.1. Participants*

Forty-seven healthy university students (31 females, mean age = 20.9 years, SD = ± 4.9) participated in the pilot study. Participants gave informed written consent before study procedures and were treated following the ethical standards of the revised Helsinki Declaration.

###### 1.1.2. *Tasks and procedure*

We randomly assigned participants to cPCT and mPCT conditions.

Trials of the two PCT versions followed the same structure (see Figure 1 in the main text – Panel A for a graphical representation): i) a fixation point was positioned at the center of a black screen, flanked by two squares on the left and right sides; ii) after an interval ranging from 200-700 ms, the outline of one of the two squares became red (cue) for 100 ms; iii) 100 ms after the cue disappeared, a small gray square (target) appeared inside the left or right square and remained on the screen until participant response for a maximum time of 2000 ms. During the task, participants were instructed to focus on the fixation point and detect the target as accurately and fast as possible, pressing the letter "F" on the Italian keyboard for targets appearing in the left square and "J" for stimuli presented to the right.

Two combinations of cue and target locations were possible: a valid or congruent condition, in which the target appeared on the same side of the bright cue, and an invalid or incongruent condition, in which the target and the cue appeared on the opposite sides. The cPCT comprised 100 trials: 80 valid trials (40 targets presented inside the right and 40 inside the left squares) and 20 invalids (10 right and 10 left). Conversely, the mPCT comprised 50 congruent (25 on the right side and 25 on the left side of the central cross) and 50 incongruent trials (25 on the right side and 25 on the left side of the central cross). In both tasks, 10 false alarm trials were added, in which no targets followed the cues. False alarms were not analyzed since participants did not have to press any key. We collected data using E-Prime Go (Psychology Software Tools, Pittsburgh, PA), an E-Prime extension allowing remote data collection. Participants received a link via email and were asked to perform the task in a quiet room, avoiding distractions. Response accuracy and reaction times (RTs) were recorded.

##### 1.1.3.*Statistical approach*

We performed statistical analyses in the R environment (R Core Team, 2022). The dependent variable accuracy was analyzed using general mixed effects models (R. H. Baayen, Davidson, & Bates, 2008), fitted using the GLMER function of the lme4 package (Bates, Maechler, Bolker, & Walker, 2015). RTs values of correct responses were analyzed according to linear mixed effects regression using the LMER procedure available in the "lme4" R package (version 1.1-5), and outliers were removed via model-criticism (2.5 SD of standardized residuals) (H. Baayen & Milin, 2010).

*Trial validity* (factorial, two levels: valid vs. invalid condition), *task* (factorial, two levels: cPCT vs. mPCT), and their interaction were entered as fixed factors in the full model. *Trial order* was also included as a continuous predictor to account for learning or fatigue effects on task performance. In the final models, fixed predictors inclusion was tested with a backward selection procedure to simplify each statistical model, thus eliminating non-significant fixed effects. The backward procedure was done manually for accuracy. Conversely, we applied for RTs the step function from the lmerTest package (version 2.0) (Kuznetsova, Brockhoff, & Christensen, 2015).

##### *1.1.4.Results*

One participant assigned to the cPCT systematically inverted the keys to perform the task, pressing "J" for targets appearing to the left and "F" to the right (mean accuracy of 5%). Another volunteer assigned to the mPCT completed the task with low accuracy (64%) compared to the rest of the group (average accuracy 96.4% ± 18.5). We removed the data collected from these two participants from subsequent analyses. A total of 45 participants was analyzed, with 23 performing the cPCT (12 females, mean age = 20.1 years, SD = ± 0.65) and 22 (19 females, mean age = 20.4 years, SD = ± 0.88) completing the mPCT.

After this procedure, we analyzed participants' accuracy on 4500 data points.

The best-fitting model was the one including the *trial order* and *trial validity* ($\chi^2_{(1)} = 49.3$, $p < .001$). The order effect highlighted that participants' accuracy improved during the task performance, showing a genuine learning effect ($\chi^2_{(1)} = 41.7$, $p < .001$). Moreover, participants were more accurate for valid (M = 99%, SD = ± 9.9) vs. invalid (M = 96.2%, SD = ± 19.2) trials ($\chi^2_{(1)} = 37.2$, $p <.001$).

Concerning RTs, analysis was run on 4411 data points. The best-fitting model was the one including the simple effect of *trial order* and the interactions between *task* and *trial validity* ($\chi^2_{\mathbf{(1)}} = 174.5$, $p <.001$). Concerning the impact of *trial order*, RTs reflected a learning effect, showing shorter responses while performing the task ($\chi^2_{\mathbf{(1)}} = 117.4$, $p <.001$). Relatively to the interaction between task and *trial validity* ($\chi^2_{\mathbf{(1)}} = 178$, $p < .001$), post-hoc analysis performed with *FDR* adjustment showed that no difference in RTs was traceable for the cPCT and mPCT valid trials (p = .911). In contrast, RTs were faster in the mPCT than the cPCT (p < .001) for invalid trials. Moreover, the well-known facilitatory effect induced by the *trial validity* was larger in the cPCT (p < .001) than in the mPCT, where the effect was still present but reduced in magnitude (p = .019) (see Figure S1).

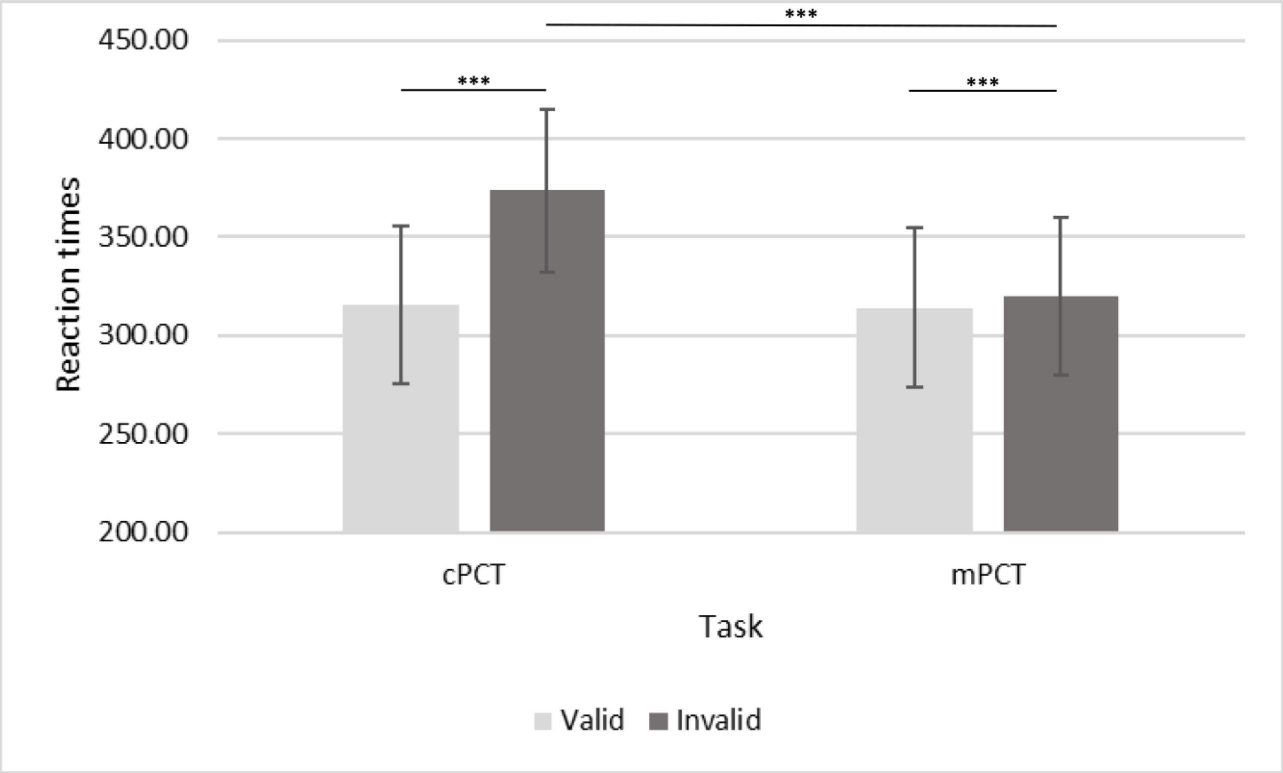


Figure S1. Results on RTs data in the cPCT and mPCT tasks.

***Table S1***. *Results of the mixed-effect analysis on accuracy in the Pilot experiment.*

| ***Parameter*** | $\chi^2$ | ***P*** | ***Removal order*** | ***Estimate*** | ***z value*** | ***p*** |
|---|---|---|---|---|---|---|
| *Intercept* | - | - | *Not removed* | 2.1709 | 10.423 | <.001 |
| *Trial validity* | - | - | *Not removed* | 1.4218 | 6.102 | <.001 |
| *Trial order* | - | - | *Not removed* | 0.0259 | 6.455 | <.001 |
| *Trial validity*task* | 2.3751 | .1233 | 1 | - | - | - |
| *Task* | 1.5967 | .2064 | 2 | - | - | - |

*Table S1 summarizes the model-simplification procedure, including the goodness-of-fit tests and their results. The rightmost part of each table reports the effects of the included variables.*

***Table S2***. *Results of the mixed-effect analysis on RTs in the Pilot experiment.*

| ***Parameter*** | $\chi^2$ | ***P*** | ***Removal order*** | ***Estimate*** | ***t value*** | ***p*** |
|---|---|---|---|---|---|---|
| *Intercept* | - | - | *Not removed* | 390.2556 | 44.263 | <.001 |
| *Task* | | | *Not removed* | -53.5258 | -4.376 | <.001 |
| *Trial validity* | - | - | *Not removed* | -57.9280 | -18.972 | <.001 |
| *Trial order* | - | - | *Not removed* | -0.264 | -10.833 | <.001 |
| *Task*trial validity* | - | - | *Not removed* | 52.187 | 13.342 | <.001 |

*Table S2 summarizes the model-simplification procedure, including the goodness-of-fit tests and their results. The rightmost part of each table reports the effects of the included variables.*

#### *1.2. Conclusions*

We built a pilot study to compare participants' performance in cPCT vs. mPCT. Crucially the main effect of task was not significant, neither on accuracy or RTs. Crucially for the purposes of our studies, the results showed that although the discrepancy between valid and invalid conditions was reduced, athe mPCT performance was comparable to that of cPCT, showing the typical advantage for valid cues.

### 2. Section B – Model selection Analyses - Experiment 1

***Table S3****. Results of the mixed-effect analysis on accuracy in the mPCT - Experiment 1.*

| ***Parameter*** | $\chi^2$ | ***P*** | ***Removal order*** | ***Estimate*** | ***z value*** | ***p*** |
|---|---|---|---|---|---|---|
| *Intercept* | - | - | *Not removed* | 3.4567 | 15.759 | <.001 |
| *Trial validity* | - | - | *Not removed* | 1.3092 | 5.368 | <.001 |
| *Trial order* | - | - | *Not removed* | 0.0097 | 2.881 | .004 |
| *TDCS condition*trial validity* | 0.9841 | .3212 | 1 | - | - | - |
| *TDCS condition* | 0.3345 | .5630 | 2 | | | |

*Table S3 summarizes the model-simplification procedure, including the goodness-of-fit tests and their results. The rightmost part of each table reports the effects of the included variables.*

***Table S4****. Results of the mixed-effect analysis on RTs in the mPCT – Experiment 1.*

| ***Parameter*** | $\chi^2$ | ***P*** | ***Removal order*** | ***Estimate*** | ***t value*** | ***p*** |
|---|---|---|---|---|---|---|
| *Intercept* | - | - | *Not removed* | 341.8422 | 75.891 | <.001 |
| *Trial validity* | - | - | *Not removed* | -0.1320 | -0.076 | .9395 |
| *TDCS condition* | - | - | *Not removed* | 4.2243 | 2.428 | .0152 |
| *Trial order* | - | - | *Not removed* | -0.1300 | -6.409 | <.001 |
| *Trial validity*tDCS condition* | - | - | *Not removed* | -8.2221 | -3.348 | <.001 |

*Table S4 summarizes the model-simplification procedure, including the goodness-of-fit tests and their results. The rightmost part of each table reports the effects of the included variables.*

***Table S5***. *Results of the mixed-effect analysis on K factor in the VWMT – Experiment 1.*

| ***Parameter*** | $\chi^2$ | ***P*** | ***Removal order*** | ***Estimate*** | ***t value*** | ***p*** |
|---|---|---|---|---|---|---|
| *Intercept* | - | - | *Not removed* | 2.7742 | 11.04 | <.001 |
| *TDCS condition*attended hemifield* | 0.0198 | .8882 | 1 | - | - | - |
| *TDCS condition* | 0.0246 | .8754 | 2 | - | - | - |
| *Attended hemifield* | 0.234 | .6286 | 3 | - | - | - |

*Table S5 summarizes the model-simplification procedure, including the goodness-of-fit tests and their results.*

### 3. Section C – Experiment 2

#### *3.1 Behavioral results – model selection*

***Table S6**. Results of the mixed-effect analysis on accuracy in the mPCT – Experiment 2.*

| ***Parameter*** | $\chi^2$ | ***P*** | ***Removal order*** | ***Estimate*** | ***z value*** | ***p*** |
|---|---|---|---|---|---|---|
| *Intercept* | - | - | *Not removed* | 4.4528 | 11.229 | <.001 |
| *Trial validity* | - | - | *Not removed* | 1.0811 | 3.658 | <.001 |
| *Trial order* | - | - | *Not removed* | -0.0104 | -2.383 | .0172 |
| *TDCS condition*trial validity* | *0.0061* | *.9376* | 1 | - | - | - |
| *TDCS condition* | *0.3194* | *.5720* | 2 | - | - | - |

*Table S6 summarizes the model-simplification procedure, including the goodness-of-fit tests and their results. The rightmost part of each table reports the effects of the included variables. The best-fitting model included trial validity and trial order as predictors.*

***Table S7**. Results of the mixed-effect analysis on RTs in the mPCT – Experiment 2.*

| ***Parameter*** | $\chi^2$ | ***P*** | ***Removal order*** | ***Estimate*** | ***t-value*** | ***p*** |
|---|---|---|---|---|---|---|
| *Intercept* | - | - | *Not removed* | 338.1327 | 42.061 | <.001 |
| *TDCS condition* | - | - | *Not removed* | -4.9932 | -3.098 | .0020 |
| *Trial order* | - | - | *Not removed* | -0.1562 | -5.882 | <.001 |
| *TDCS condition*trial validity* | *0.6456* | *.4217* | 1 | - | - | - |
| *Trial validity* | *0.1195* | *.7296* | 2 | | | |

*Table S7 summarizes the model-simplification procedure, including the goodness-of-fit tests and their results. The rightmost part of each table reports the effects of the included variables. The best-fitting model included the tDCS condition and the trial order as predictors.*

***Table S8**. Results of the mixed-effect analysis on VWMT K-factor – Experiment 2.*

| ***Parameter*** | $\chi^2$ | ***P*** | ***Removal order*** | ***Estimate*** | ***t value*** | ***p*** |
|---|---|---|---|---|---|---|
| *Intercept* | - | - | *Not removed* | 3.0607 | 9.47 | <.001 |
| *TDCS condition*attended hemifield* | 0.6753 | .4112 | 1 | - | - | - |
| *TDCS condition* | 0.0057 | .9397 | 2 | - | - | - |
| *Attended hemifield* | 0.0013 | .9718 | 3 | - | - | - |

*Table S8 summarizes the model-simplification procedure, including the goodness-of-fit tests and their results. The rightmost part of each table reports the effects of the included variables. The best-fitting model was the one including the tDCS condition as predictor.*

### *3.2* TMS-EEG Analyses

#### *3.2.1 Analysis approach for TMS-EEG data*

We performed three analyses on our EEG dataset for c-tDCS and Sham conditions. Two analyses were at the sensor level and one at the level of cortical sources.

#### *3.2.1 GMFP and LMFP*

At the sensors level, mirroring the analysis of our previous studies (Romero Lauro et al., 2016, 2014; Varoli et al., 2018), we computed the GMFP and the LMFP values for each experimental condition (pre-and post-tDCS) and within three-time windows (0–50, 50–100, and 100–150 ms after the TMS pulse). In a second moment, following the cluster analysis results, we decided to extend the time window up to 300 ms to check for the possibility of late effects. Therefore, we added three more-time windows from 150 to 300 ms after the TMS pulse (150-200, 200-250, and 250-300 ms). We computed GMFP considering signals derived from all 60 channels. Instead, we calculated LMFP in four electrode clusters, each made of 4 electrodes, located bilaterally in the frontal and parietal lobes. The left parietal cluster (Cluster 1: CP1, CP3, P1, and P3) corresponded to the TMS hotspot and the right one (Cluster 2: CP2, CP4, P2, and P4) to the tDCS cathode position. The two frontal clusters corresponded to the areas structurally and functionally connected to the parietal ones: the left frontal (Cluster 3: F1, F5, FC1, and FC3) and the right frontal clusters (Cluster 4: F2, F6, FC2, and FC6).

Then, the GMFP and LMFP data were submitted to a series of linear mixed models using the "lme4" package (version 1.1 5) (Bates et al., 2015) built for the R statistical software (R Core Team, 2022). On the same GMFP and LMFP data, we performed a further test with a Bayesian ANOVA to test for the null hypothesis (Etz, Gronau, Dablander, Edelsbrunner, & Baribault, 2018; Rouder, Speckman, Sun, Morey, & Iverson, 2009) through the Bayesian Analysis of Variance (ANOVA) using "JASP" software environment (version 0.8.2.0) (JASP Team, 2017).

#### 3.2.2 *Results of sensor analysis for c- and sham tDCS*

##### *3.2.2.1 C-tDCS*

*Global Mean Field Power – GMFP:* in the model used to describe the GMFP values, the condition's main effect was not significant. This result means that GMFP did not change when TEPs were recorded before and after c-tDCS. Specifically, the model did not include the main effect of the *condition* in the 0-50 ms ($\chi^2_{(2)}$ = 0.12, p = .732), in the 50-100 ms ($\chi^2_{(2)}$ = 1.04, p = .324) and in the 100-150 ms ($\chi^2_{(2)}$ = 0.04, p = .852) time-windows (see Figure S2). Bayesian repeated ANOVA analysis provided only anecdotal evidence favoring the null hypothesis (see Table S9 for details on Bayesian results).

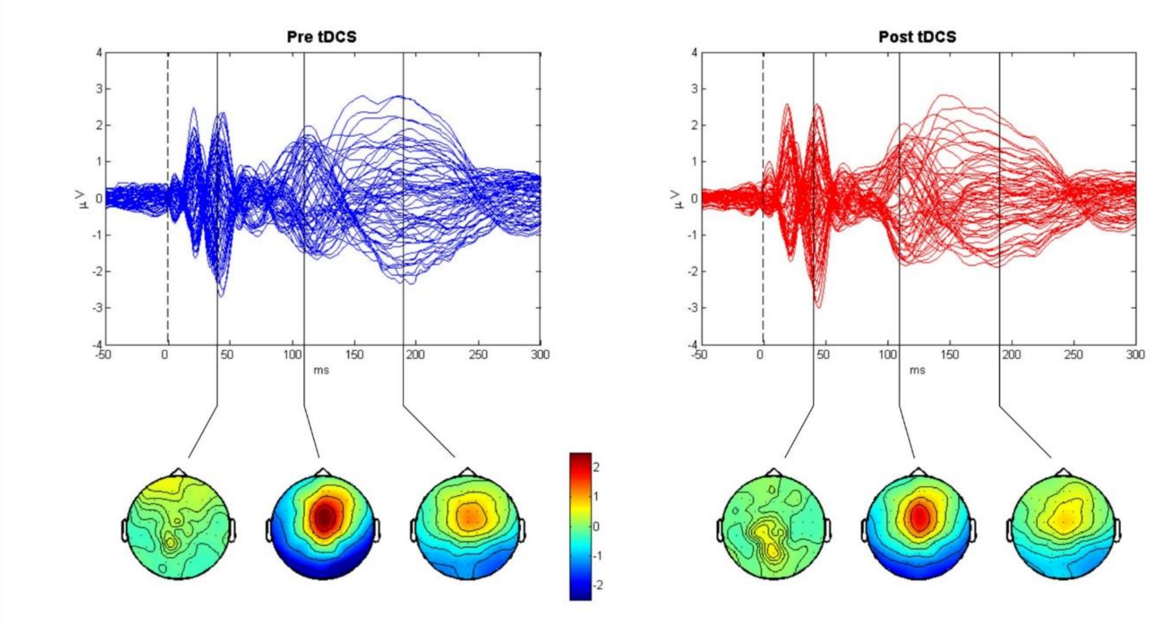


Figure S2: Results of the GMFP for the c-tDCS condition. In the upper bound is depicted the butterfly plot of the 60 channels' TEPs for the two conditions: Pre-tDCS in blue and Post-tDCS in red. The lower row represents the mean topographies computed in correspondence with the local maxima for the following time windows: 0-50 ms, 100-150 ms, and 150-200 ms.

*Local Mean Field Power – LMFP:* Analyses performed on LMFP values did not indicate any effect of c-tDCS in any of the considered clusters of electrodes. Below, the results for the four electrode clusters are given in detail.

Concerning Cluster1, the cluster corresponding to the left PPC, LRT indicated not to include *condition* in models for any time window (0-50 ms: $\chi^2_{(2)}$ = 2*10-4; p = .989; 50-100 ms: $\chi^2_{(2)}$ = 0.09; p = .765; 100-150 ms: $\chi^2_{(2)}$ = 0.06; p = .813).

Considering Cluster2, the cluster under the tDCS cathode, the model on LMFP values did not include the main effect *condition* in the first ($\chi^2_{(2)}$ = 0.81; p = .383) second ($\chi^2_{(2)}$ = 0.09; p = .766) and third ($\chi^2_{(2)}$ = 0.63; p = .431) time-window.

Similarly, in Cluster3, the left frontal cluster (F1, F5, FC1, and FC3), the analysis did not include the main effect of *condition* in models for any time window (0-50 ms: $\chi^2_{(2)}$ = 0.67; p = 0.407; 50-100 ms: $\chi^2_{(2)}$ = 0.99; p = .314; 100-150 ms: $\chi^2_{(2)}$ = 0.24; p = .629).

Lastly, on Cluster4, the right frontal cluster (F2, F6, FC2, and FC6), the final model did not include the main effect of *condition* in the 0-50 ms ($\chi^2_{(2)}$ = 0.12; p = .739), nor in the 50-100 ms ($\chi^2_{(2)}$ = 1.64; p = .200), or in the 100-150 ms ($\chi^2_{(2)}$ = 0.06; p = .816) time windows.

For all four clusters of interest, the Bayesian analyses revealed anecdotal or moderate evidence in favor of the null hypothesis (see Table S9 for details on Bayesian results).

##### *3.2.2.2 Sham tDCS*

*Global Mean Field Power – GMFP:* As expected, sham stimulation did not modulate global indices of cortical excitability. In particular, the final model run on GMFP did not include the main effect of the *condition* in any time window (0-50 ms: $\chi^2_{(2)}$ = 0.17; p = .688; 50-100 ms: $\chi^2_{(2)}$ = 0.14; p = .713; 100-150 ms: $\chi^2_{(2)}$ = 0.57; p = .453) (see Figure S3).

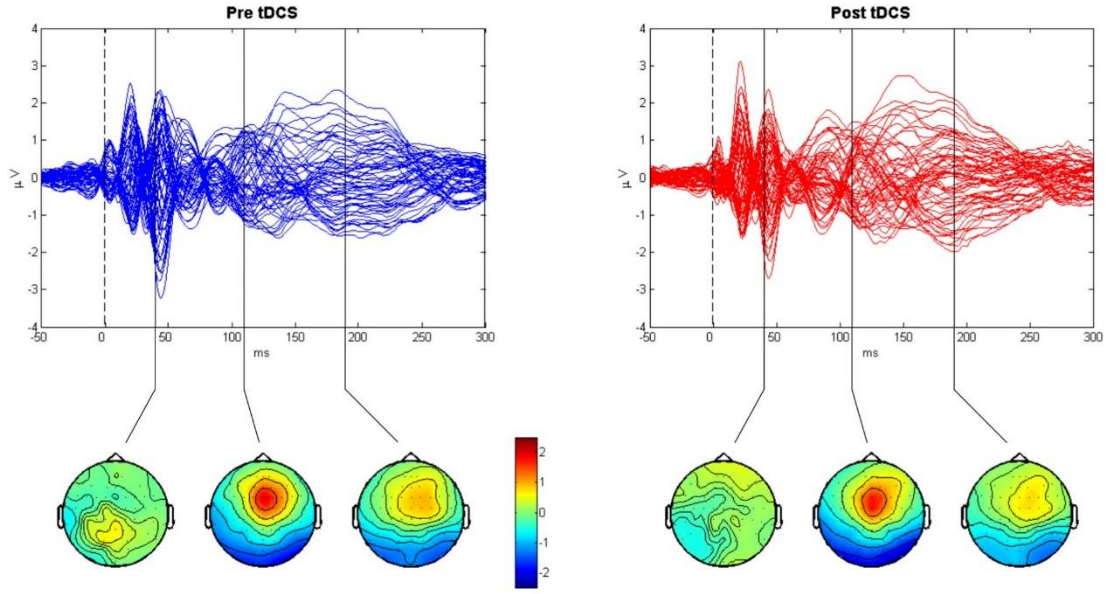

Figure S3: Results of the GMFP for the sham condition. In the upper bound is depicted the butterfly plot of the 60 channels' TEPs for the two conditions: Pre-tDCS in blue and Post-tDCS in red. The lower row represents the mean topographies computed in correspondence with the local maxima for the following time windows: 0-50 ms, 100-150 ms, and 150-200 ms.

*Local Mean Field Power – LMFP:* Analyses run on LMFP confirmed the lack of any effect of sham tDCS in modulating local cortical excitability in any of the considered clusters of electrodes.

In Cluster1, LRT values were non-significant for the first ($\chi^2_{(2)} = 0.15$; p = .696), the second ($\chi^2_{(2)} = 0.56$; p = .463) and the third ($\chi^2_{(2)} = 1.14$; p = .292) time-window. The same holds for Cluster 2. LRT values were not significant for the factor *condition* in any time window (0-50 ms: $\chi^2_{(2)} = 0.002$; p = .961; 50-100 ms; $\chi^2_{(2)} = 0.02$; p = .897; 100-150 ms: $\chi^2_{(2)} = 1.14$; p = .291). Similarly, in Cluster3, LRT indicated not to include factor *condition* for any time window (0-50 ms: $\chi^2_{(2)} = 0.12$; p = .736; 50-100 ms: $\chi^2_{(2)} = 0.96$; p = .333; 100-150 ms: $\chi^2_{(2)} = 1.59$; p = .226) in the final model. Finally, also for Cluster 4, LRT values were non-significant for the first ($\chi^2_{(2)} = 0.02$; p = .896), the second ($\chi^2_{(2)} = 1.13$; p = .293), and the third ($\chi^2_{(2)} = 0.07$; p = .796) time-window.

For the sham stimulation condition, the results of the Bayesian analysis also revealed anecdotal or moderate evidence in favor of the null hypothesis, with BF values between 2.0 and 3.0 (Supplementary materials - section C, Table S9).

#### *3.2.3 Results of Bayesian ANOVA*

##### *3.2.3.1 Cathodal stimulation condition – Bayesian Factors*

***Table S9**. Results of the Bayesian ANOVA in the GMFP and LMFP data from the c-tDCS condition.*

| ***Time window (ms)*** | ***GMFP*** | ***LMFP*** | | | |
|---|---|---|---|---|---|
| | | Cluster1 | Cluster2 | Cluster3 | Cluster4 |
| *0 - 50* | *2.8* | *2.1* | *2.1* | *2.5* | *2.5* |
| *50 - 100* | *2.3* | *2.9* | *3.0* | *2.7* | *2.6* |
| *100 – 150* | *3.0* | *3.0* | *2.4* | *2.6* | *2.6* |

*Table S9 summarizes the value of the Bayesian Factor ($BF_{01}$) for each time window. The table shows the results for both GMFP and LMFP data. $BF_{01}$ values ranged between 2.1 and 3.0, revealing anecdotal (range 1 - 3) and moderate (range 3 – 10) evidence in favor of the null hypothesis.*

*3.2.3.2 Sham stimulation condition – Bayesian Factors*

***Table S10.*** *Results of the Bayesian ANOVA in the GMFP and LMFP data from the sham tDCS condition.*

| ***Time window (ms)*** | ***GMFP*** | ***LMFP*** | | | |
|---|---|---|---|---|---|
| | | Cluster 1 | Cluster 2 | Cluster 3 | Cluster 4 |
| *0 - 50* | *3.0* | *2.7* | *3.0* | *2.9* | *2.5* |
| *50 - 100* | *2.9* | *2.5* | *1.3* | *2.4* | *2.7* |
| *100 – 150* | *2.9* | *2.0* | *2.3* | *2.3* | *2.6* |

*Table S10 summarizes the value of the Bayesian Factor ($BF_{01}$) for each time window. The table shows the results for both GMFP and LMFP data. $BF_{01}$ values ranged between 1.3 and 3.0, revealing anecdotal (range 1 - 3) and moderate (range 3 – 10) evidence favoring the null hypothesis.*

*3.2.4. Correlations between the neurophysiological and behavioral data*

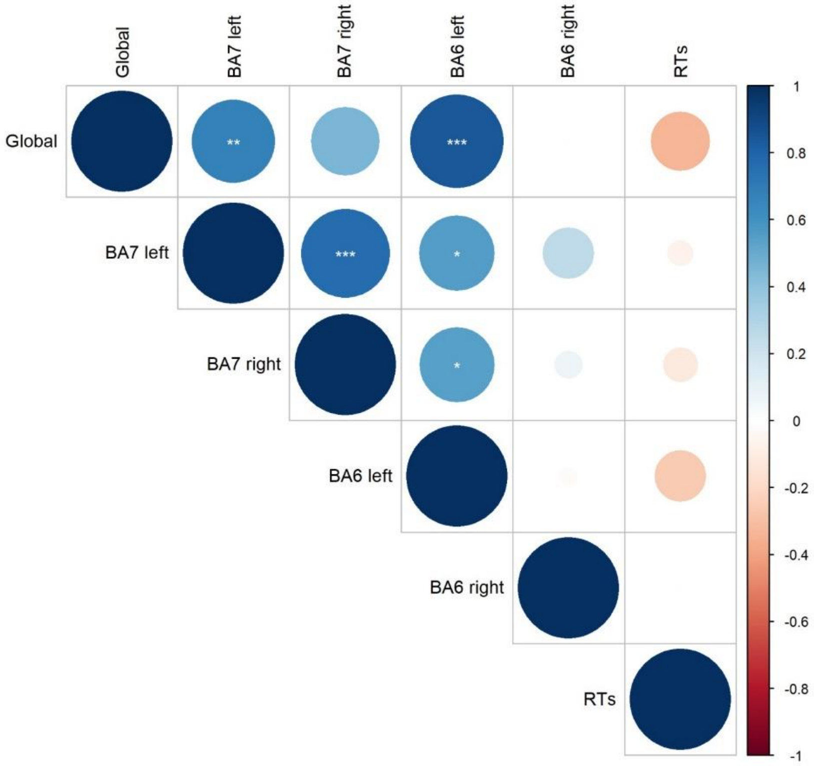


Figure S4: The correlogram or correlation matrix depicts the relationship between each pair of variables. Positive correlations are blue colored, and negative ones are in red. The color intensity and dots size are proportional to the correlation coefficients, and the asterisks inside the dots represent the statistical significance (* p < .05, ** p < .01, *** p < .001).

As highlighted by the correlation matrix, no significant correlations emerged between the neurophysiological and behavioral data. Positive correlations emerged between SCD of different regions.
